# A variational method for efficient estimation of diffusion and free-energy profiles along collective variables

**DOI:** 10.64898/2026.01.01.697292

**Authors:** Anže Hubman, Franci Merzel

## Abstract

An efficient variational method is presented for estimating the diffusion coefficients and free-energy profiles along selected collective variables from projected molecular dynamics trajectories under both equilibrium and nonequilibrium conditions. The method is based on the assumption that the short-time transition probability density of the coordinate moves can be approximated by a Gaussian form. Defining a loss function as the sum of Kullback-Leibler divergences between the analytical short-time propagators of an overdamped Langevin model and those estimated directly from the projected trajectories maximises the agreement between the two and allows for its analytic evaluation. To efficiently minimise this loss function by varying diffusion and free-energy profiles along collective variables, we use an adaptive Monte Carlo scheme. The method is applied to two model systems exhibiting diffusive dynamics, as well as to water diffusion across the interface of a biomolecular condensate, demonstrating its robustness and accuracy.

## 1 Introduction

Molecular dynamics (MD) simulations have advanced to the point where they can be used to quantitatively model complex phenomena, including protein conformational dynamics [1, 2], self-assembly of biomolecules in solution [3], permeation of water and ions through biological membranes [4,5], chemical reactions on catalytic surfaces [6], and phase transitions in crystalline materials [7, 8]. In MD simulations, the equations of motion are integrated numerically, which produces trajectories in the 6*N*-dimensional phase space (*N* being the number of particles) that describe the time evolution of particle positions and momenta. A posteriori analysis of the simulated trajectories provides access to the structural, dynamical, and kinetic properties of the system [9].

Because MD trajectories are inherently high-dimensional, analyzing them typically requires dimensionality reduction [10]. A low-dimensional effective description of the system can be constructed by projecting the trajectories onto a small set of collective variables, which are functions of the generalized coordinates that characterize key states of the system, such as reactants, products, and transition states [11]. The construction of optimal collective variables is problem-specific and is guided by physical intuition [12] or data-driven methods [13, 14]. The chosen set of collective variables, **q** = (*q*_1_, …, *q*_*N*_), uniquely defines the associated free-energy surface *F* (**q**), which can be determined accurately and efficiently using enhanced sampling techniques [15–17]. Formally, the projected dynamics follows the non-Markovian generalized Langevin equation (GLE), which is applicable over arbitrary timescales [18–20]. In addition to the free-energy surface, the GLE depends on the memory kernel that encodes memory effects responsible for non-Markovian dynamics. Despite significant progress, accurate parameterization of the memory kernel remains numerically challenging and requires long trajectories sampled at high temporal resolution [21–30]. However, for timescales longer than the characteristic memory time, non-Markovian effects can be neglected. By assuming that the instantaneous velocity 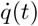 follows the equilibrium distribution, the time evolution of *q* at an inverse temperature *β* is given by the Markovian overdamped Langevin equation [31]:

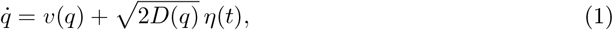

in which the generalized drift *v*(*q*) is expressed as the sum of the thermodynamic force due to the free-energy profile *F* (*q*) and the drift originating from position-dependent diffusion coefficient

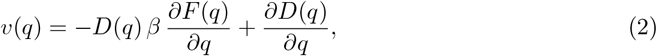

where *η*(*t*) denotes Gaussian white noise, which satisfies ⟨*η*(*t*)⟩ = 0 and ⟨*η*(*t*)*η*(*t*′)⟩ = *δ*(*t* − *t*′). Overdamped Langevin models have been successfully used to describe the kinetics of protein folding [32,33], conformational transitions of proteins [34], looping of a polymer chain in solution [35], nucleation of vapor bubbles [36], and the dynamics of self-association between pairs of fullerenes in water [37]. To obtain an accurate description of the kinetics, it is often necessary for the diffusion coefficient to depend on the collective variable. However, extracting *D*(*q*) from the time series of *q* is challenging.

The primary goal of many existing methods is to construct a likelihood function that quantifies the probability of observing a specific trajectory in the collective-variable space, given the model parameters *F* (*q*) and *D*(*q*). To construct the likelihood, an explicit expression for the propagator *p*(*q*′, *τ* |*q*, 0) is required. The latter expresses the conditional probability of observing a system in a state *q*′ at time *τ*, given that it was in a state *q* at time *t* = 0. Building on the work of Bicout and Szabo [38], Hummer [39] discretized the collective-variable space and expressed the local propagators of an overdamped Langevin model in terms of the lag-time *τ* and a rate matrix **R** that describes transitions between adjacent cells and encodes information on *F* (*q*) and *D*(*q*). In this framework, the likelihood depends on the number of transitions that occur between neighboring cells within the time interval *τ* and on the rate matrix **R**, whose elements are treated as parameters. The elements of **R** that best reproduce the observed trajectory along *q* are obtained by Bayesian inference or by maximizing the likelihood, yielding self-consistent estimates of *F* (*q*) and *D*(*q*). A key requirement of this approach is that *τ* be chosen such that a sufficient number of nearest-neighbor transitions are observed.

Analytical expressions for the short-time propagators can also be used [37,40]. In this case, *F* (*q*) and *D*(*q*) are directly varied to maximize the likelihood of the observed trajectory. The choice of *τ* is crucial in this method, since it must be short enough for the analytical approximation to hold but long enough for the Markovian assumption to remain valid. Although this approach has strong statistical foundations, the evaluation of the likelihood function at every optimization step becomes computationally expensive for long trajectories. A related approach is to perform this maximization analytically with respect to *v*(*q*) and *D*(*q*), which gives the first and second Kramers-Moyal coefficients that link the conditional averages and variances of the displacements Δ*q* = *q*(*t*+*τ*)−*q*(*t*) to the drift and diffusion terms in Eq. 1 [35,41]. While this avoids variational optimization, it requires verification that the reconstructed *F* (*q*) from *v*(*q*) and *D*(*q*) is consistent with the true free-energy profile.

Alternative methods avoid explicit likelihood construction and are generally less sensitive to the choice of *τ*. However, they are typically more computationally demanding. Liu *et al*. introduced a dual-simulation approach, in which Langevin dynamics simulations are performed with a known free-energy profile, and the diffusion coefficient is adjusted until the simulated survival probability matches that from MD [42]. Woolf and Roux estimated *F* (*q*) and *D*(*q*) from a series of locally restrained simulations [43], while Pérez-Villa and Pietrucci optimized the free-energy, friction, and mass profiles by matching the time-dependent distribution 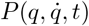 obtained from a few hundred short MD trajectories initiated at the transition state to that generated by Langevin dynamics, thus capturing both underdamped and overdamped regimes [44].

In this work, we present a simple and computationally efficient variational approach for estimating diffusion coefficients and, optionally, free-energy profiles along selected collective variables from projected MD trajectories. The method is based on a loss function designed to maximize the agreement between the analytical short-time propagators of an overdamped Langevin model, defined by a given *F* (*q*) and *D*(*q*), and the corresponding propagators estimated directly from the projected trajectories. We show that an adaptive Monte Carlo minimization scheme arises naturally from the construction of the loss function. The robustness of the approach is demonstrated on two model systems exhibiting diffusive dynamics under both equilibrium and non-equilibrium conditions, and by analyzing water diffusion between the dense and dilute phases of a biomolecular condensate.

## 2 Methods

### a) Variational optimization

We outline a variational procedure for determining the position-dependent diffusion coefficient *D*(*q*) along a single collective variable *q*. The method can be extended to multiple dimensions if substantially more data is available. Unless stated otherwise, the free-energy profile is assumed to have been estimated independently.

Consider a discrete time series *q*(*t*_*i*_) = *q*_*i*_, sampled at temporal resolution *t*_*i*+1_ − *t*_*i*_ = Δ*t*. The lag-time *τ* is defined as *τ* = *k*Δ*t*, where *k* is a positive integer. For sufficiently large *τ*, we assume that the time evolution of *q* can be approximated as Markovian and described by an overdamped Langevin equation parameterized in terms of *F* (*q*) and *D*(*q*). Under this assumption, the conditional probability of making a step Δ*q* over a time interval *τ* from the initial position *q*, (i. e. the propagator) has a simple Gaussian form [35, 37]:

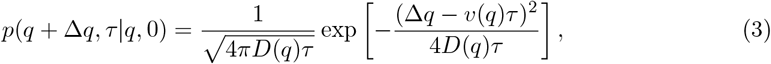

where *v*(*q*) is given by Eq. 2. This expression is valid if *F* (*q*) and *D*(*q*) are approximately constant over the displacement Δ*q* which occurs in time *τ*.

The propagators, which contain information about *F* (*q*) and *D*(*q*), can be estimated directly from the recorded time series and are referred to as empirical in this work. To compute these empirical propagators, we assume that 0 ≤ *q*(*t*) *< L* and discretize the collective-variable space into *N* bins of equal width *δ*= *L/N*. Along the observed trajectory, we collect the displacements Δ*q* = *q*(*t* + *τ*) − *q*(*t*) and assign the displacement to the *j*-th bin if *jδ*≤ *q*(*t*) *<* (*j* + 1)*δ*. For each bin, the empirical propagator is determined by computing the sample mean 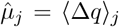 and variance 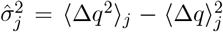, which fully specify the propagator under the assumption that the displacement distribution is Gaussian.

To obtain a self-consistent estimate of *D*(*q*) (and optionally *F* (*q*)), we require that for a given choice of free-energy and diffusion profile, the analytical propagators (*p*_*j*_) match the empirical ones 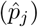 as closely as possible. More specifically, we introduce a loss function ℒ as the sum of Kullback-Leibler (KL) divergences D_KL_ between *p*_*j*_ and 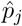

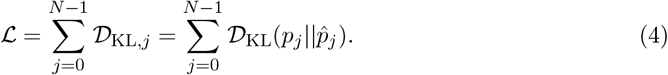

Additional regularization terms that impose smoothnes on *F* (*q*) and *D*(*q*) can be added to the loss [39]. The KL-divergence is defined as:

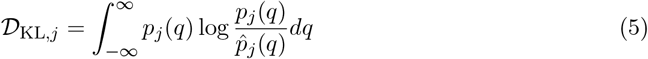

and vanishes if 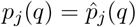 For two normal distributions *p*_*j*_ = 𝒩 (*µ, σ*) and 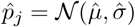, the integral in Eq. 5 can be evaluated analytically, yielding

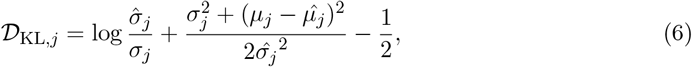

where, according to Eqs. 2 and 3, *µ* = *v*(*q*_*j*_)*τ* and *σ*^2^ = 2*D*(*q*_*j*_)*τ*.

The loss, which is a functional of both *F* (*q*) and *D*(*q*), is minimized using a Monte Carlo (MC) scheme. This approach is preferred over gradient-based algorithms because it does not require evaluation of the gradient of the loss function and allows the strict positivity of *D*(*q*) to be easily enforced. By representing *F* (*q*) and *D*(*q*) on a discrete grid of *N* points, the functional minimization is transformed into a multivariate optimization problem over the corresponding grid values. Each MC step consists of proposing a small perturbation of the selected grid value, which is then accepted or rejected based on the Metropolis criterion, treating the loss as an effective energy function. Importantly, the computational cost of evaluating the loss scales as *N*, while for the approaches based on maximum-likelihood, the cost instead scales with the length of the trajectory.

The minimization can be performed very efficiently by invoking several assumptions. Firstly, the free-energy is kept fixed at its initial estimate. This choice is motivated by the fact that *F* (*q*) can be estimated relatively accurately using standard histogramming techniques or more advanced methods when necessary. Secondly, because only the diffusion profile *D*(*q*) is varied during minimization, the locations of the proposed MC moves can be *optimally* selected. This is achieved by converting the set of KL-divergences 𝒟 _KL,*j*_ from a previous optimization step into a discrete probability distribution *P* defined as:

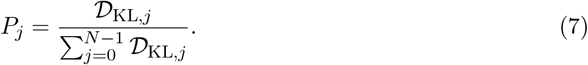

New locations for the proposed MC moves are sampled from the distribution *P* rather than being chosen randomly. This strategy allows the algorithm to adaptively focus on regions of *D*(*q*) that contribute most strongly to the loss. A limitation of keeping the free-energy profile fixed is that any error in *F* (*q*) propagates to the estimation of *D*(*q*). This issue can be mitigated by performing a few additional optimization steps in which both *F* (*q*) and *D*(*q*) are allowed to vary.

The accuracy of the method critically depends on the validity of the analytical expression for the propagator at the chosen value of *τ*. When *τ* is too small, non-Markovian effects become significant, whereas for sufficiently large *τ*, Eq. 3 is no longer valid. To enable the use of larger values of *τ*, more accurate approximations of the propagator can be employed. For example, in Ref. [37], the authors used a Gaussian-form propagator that additionally incorporates higher-order derivatives of *F* (*q*) and *D*(*q*). Alternatively, a more computationally demanding but stable approach for large *τ* consists of performing repeated Langevin dynamics simulations on *F* (*q*) and *D*(*q*) at each optimization step, constructing propagators from both MD and Langevin trajectories as discrete histograms, and evaluating Eq. 5 numerically.

### b) Diffusion on a periodic domain

We tested the method on a one-dimensional model system undergoing diffusive dynamics on a periodic domain [39]. The free-energy and diffusion profiles were defined as

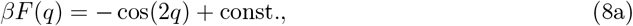

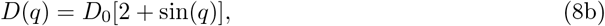

where 0 ≤ *q <* 2*π* and *D*_0_ = 0.1 rad^2^ ps^−1^. The trajectory used for the analysis was generated by integrating the overdamped Langevin equation for 100 ns with a timestep of 0.01 ps using the Euler–Maruyama integrator.

The diffusion profile was reconstructed using *N* = 24 local propagators, with a lag-time of *τ* = 5Δ*t*. The minimization was initiated from a uniform diffusion profile with *D*(*q*) = 0.2 rad^2^ps^−1^. The analytical expression for the free-energy was used to separate the estimation of *D*(*q*) from the inaccuracies of *F* (*q*). Statistical uncertainties were quantified by performing ten independent simulations and estimating the diffusion profile for each realization.

### c) Coexistence simulation of FUS-LCD

To demonstrate the applicability of our method to a realistic system, we simulated phase co-existence between the dilute and dense phases of the low-complexity domain of the Fused in Sarcoma protein (FUS-LCD) using the coarse-grained Martini3-IDP force field [45]. An equilibrated configuration consisting of 36 protein chains was taken from the work of Wang *et al*. [45] and placed at the center of an elongated simulation box with dimensions 12 nm × 12 nm × 50 nm. The system was solvated with 44,504 water molecules (see Fig. 1). A salt concentration of 0.1 M was achieved by adding 488 Na^+^ and 416 Cl^−^ ions. After a short equilibration period in the NPT ensemble, the system was simulated for 150 ns in the NVT ensemble at *T* = 300 K using GROMACS 2025.2 [46]. Trajectories were recorded every 4 ps. The default parameters recommended for the Martini3-IDP force field and periodic boundary conditions were used.

**Figure 1:**
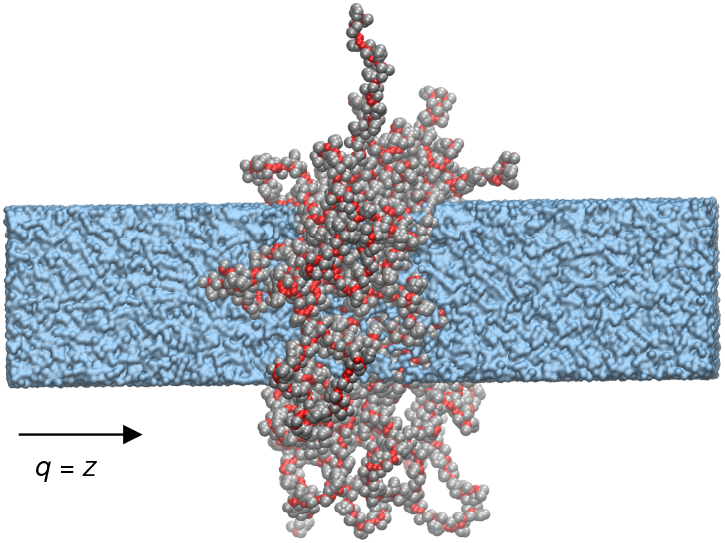
Coexistence between the dilute and dense phases of the FUS-LCD condensate simulated using the slab method. Because of the relatively short simulation time, no exchange of proteins between the two phases was observed. Backbone and side-chain beads are shown in red and gray, respectively, and the solvent is depicted in blue. Sodium and chloride ions are omitted.

We focused on the position-dependent diffusion coefficient of water molecules along the *z* direction, i.e., perpendicular to the interface. The collective variable *q* was defined as the *z* coordinate of water molecules. The free-energy profile was obtained from

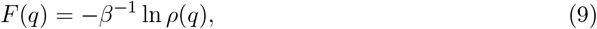

where *ρ*(*q*) is the normalized histogram of *q* obtained from the MD trajectory. The position-dependent diffusion coefficient, *D*(*q*), was estimated using *N* = 30 local propagators and a lag-time of *τ* = 50Δ*t*. Minimizations were initiated from a uniform diffusion profile *D*(*q*) = 0.2 Å^2^ps^−1^. Statistical uncertainties were quantified by dividing the trajectory into three 50-ns blocks, with the free-energy and diffusion profiles estimated independently for each block.

To assess the accuracy of the estimated *F* (*q*) and *D*(*q*), the distributions of first-passage times obtained from molecular dynamics trajectories were compared with those predicted by a Langevin model. Assuming that the dense phase is centered at *q* = 0, a first-passage event was defined as the time required for a molecule initially located within *q <* ±*L/*10 to exit the dense phase, whose boundaries were set at *q* = ±*L/*4, where *L* is the length of the simulation box along the *z* direction. First-passage times were extracted from the MD trajectories by analyzing the motion of water molecules along the *z* coordinate. Langevin dynamics simulations, with initial conditions randomly sampled near *q* = 0, were performed using the estimated *F* (*q*) and *D*(*q*), and the resulting trajectories were analyzed identically to the MD data.

### d) Double-well potential

In the third example, we simulated diffusive dynamics in a double-well potential. The free-energy and diffusion profiles are given by

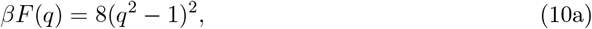

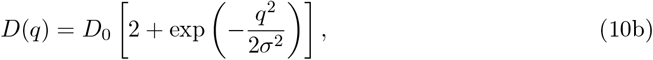

where *D*_0_ = 0.01 ps^−1^ and *σ* = 0.3. In contrast to the first model system, where the collective-variable space was easily explored, this potential contains a much higher free-energy barrier of 8*k*_B_*T*, making ergodic sampling difficult. To explore the region between the two minima, we followed the approach of Ref. [37] and relaxed 400 short trajectories from the top of the barrier. The trajectories were generated using the Euler–Maruyama integrator with a timestep of 10^−3^ ps and propagated for 2 × 10^4^ steps, sufficient to observe relaxation into one of the minima.

This example was designed to test whether a relatively small set of non-equilibrium trajectories is sufficient to recover *F* (*q*) and *D*(*q*) with our method. Since the free-energy profile cannot be estimated by histogramming, the minimization was initialized with *F* (*q*) = 0 and *D*(*q*) = 0.02 ps^−1^. In each Monte Carlo step, whether to perturb *F* (*q*) or *D*(*q*) was decided randomly. We used *N* = 30 local propagators and a lag time of *τ* = 10Δ*t*. Statistical uncertainties were quantified by performing ten independent sets of short simulations and estimating the diffusion profile (and the free-energy) for each set.

## 3 Results and Discussion

We assess the performance of the proposed method using three representative examples introduced in the previous Section. In all examples, the error bars correspond to ±*σ*, where *σ* is the sample variance.

First, we examine the diffusion on a periodic domain. As shown in Figs. 2a and 2b, the method accurately recovers the true diffusion profile, with only minor discrepancies in the region where both *F* (*q*) and *D*(*q*) exhibit local maxima. The reconstructed profile is independent of the strategy used to propose the locations of new MC moves. However, employing an adaptive (biased) proposal scheme reduces the number of optimization steps by approximately a factor of 1.8. As illustrated in Fig. 2c, the algorithm correctly identifies regions where the deviation between the initial guess and the true diffusion profile is the largest. Moreover, the local KL-divergences, used to construct weights for proposing new MC move locations, become increasingly lower and uniform as optimization proceeds.

**Figure 2:**
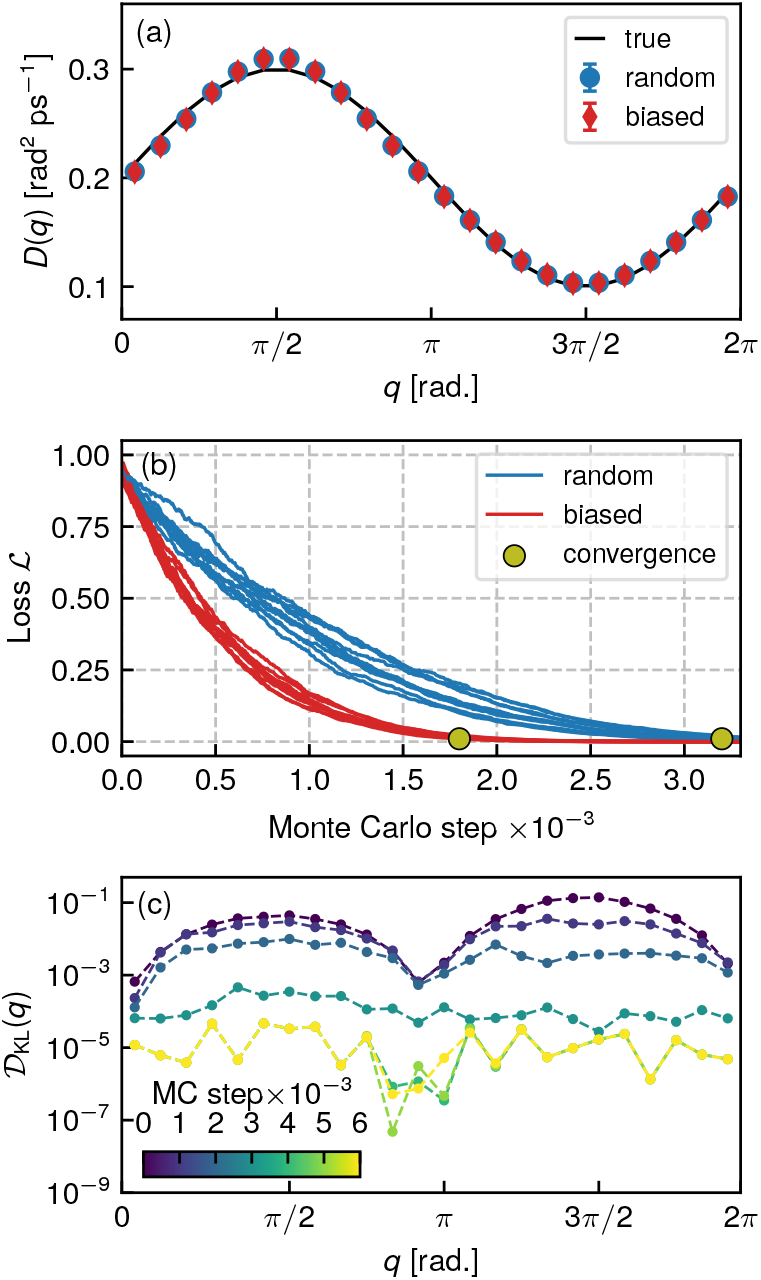
(a) Diffusion profiles of a model system moving on a periodic domain reconstructed using uniformly distributed (blue symbols) and biased (red symbols) Monte Carlo move locations. (b) Comparison of the convergence rates for random and biased Monte Carlo moves, with each line representing the realization of a minimization procedure. (c) An example evolution of local KL-divergences during Monte Carlo minimization of the loss.

In the second example, we examine the diffusion of water across the interface of a biomolecular condensate. Figure 3a shows the free-energy profile along the *z*-direction, which exhibits a small barrier of approximately 0.5 *k*_B_*T*. This is consistent with the rapid exchange of water molecules between the dilute and dense phases.

**Figure 3:**
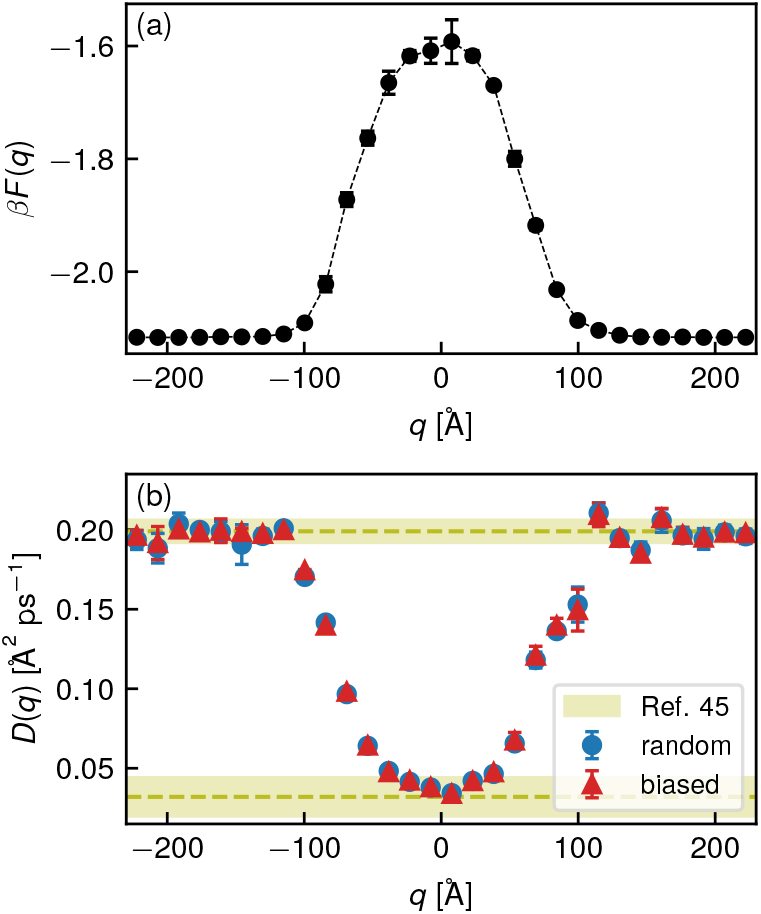
(a) Free-energy profile along the *z*-coordinate of water molecules estimated using the standard histogramming technique. (b) Corresponding position-dependent diffusion profiles computed using random (blue symbols) and biased (red symbols) Monte Carlo proposals. Reference diffusion coefficients for Martini 3 water (yellow) in the dilute and dense phases of FUS-LCD were taken from Wang et al. [45], who computed these values using a local mean-squared displacement approach.

Figure 3b presents the corresponding diffusion profile along the *z*-direction. The final estimates of *D*(*q*) are independent of the specific Monte Carlo (MC) move proposal scheme. We obtain *D*_wat_ = 0.198 ± 0.004 Å^2^/ps in the dilute phase and *D*_wat_ ≈ 0.034 ± 0.001 Å^2^/ps in the dense phase. These values are in excellent agreement with those of Wang *et al*. [45], who reported *D*_wat_ = 0.199±0.007 Å^2^/ps and 0.032±0.012 Å^2^/ps for the dilute and dense phases, respectively. Their analysis was based on local mean-squared displacements, which cannot be applied in regions where the curvature of the free-energy is nonzero, whereas our method remains valid throughout the entire interfacial region.

Figure 4 compares the efficiency of optimization schemes differing in how new MC moves are proposed. Although both approaches converge to the same minimum loss, biased proposals require roughly three times fewer optimization steps. The exact performance gain depends on the number of bins used to discretize *D*(*q*) and on the fraction of bins where *D*(*q*) can be given a good initial guess. This makes our method particularly suitable for interfacial systems, where the diffusion coefficient can be accurately determined far from the interface, allowing the algorithm to focus optimization on the interfacial region.

**Figure 4:**
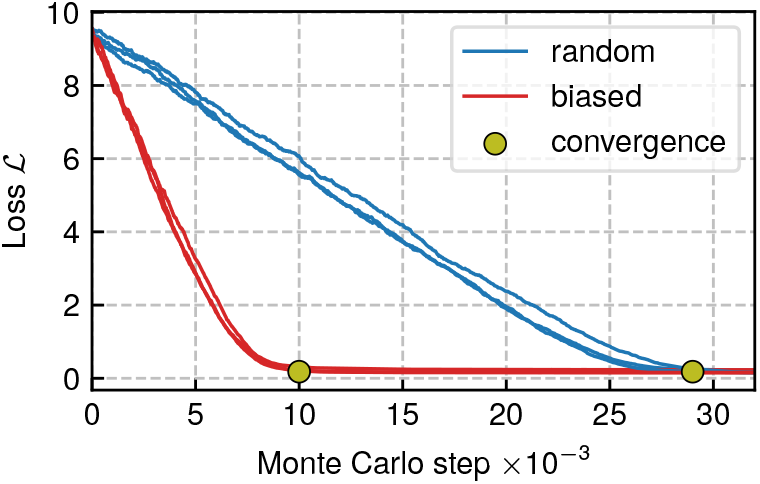
Comparison of convergence rates for random and biased Monte Carlo proposal schemes applied to water diffusion across an interface of a FUS-LCD condensate. Each line represents a separate realization of the minimization procedure.

Finally, Figure 5 compares first-passage-time (FPT) distributions obtained directly from MD simulations with those predicted by the Langevin model. This serves as a self-consistency test, since both *F* (*q*) and *D*(*q*) influence FPT distributions when the free-energy barrier is small. The Langevin model closely reproduces the MD results as evidenced by a close overlap of both FPT distributions. Furthermore, the mean first-passage time from MD simulations is ⟨*τ*_FP_⟩ = 41 ± 3 ns, while the Langevin model yields ⟨*τ*_FP_⟩ = 37.7 ± 0.7 ns.

**Figure 5:**
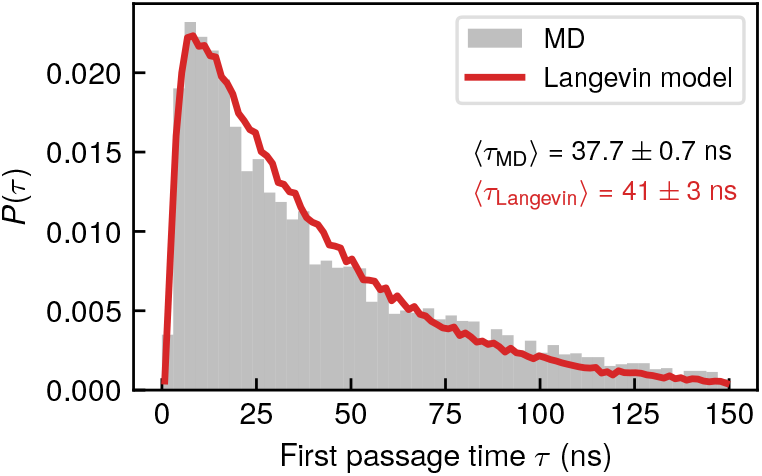
Comparison between first passage time distributions of water molecules leaving the dense phase computed from molecular dynamics simulations and those predicted by a Langevin model.

The third example differs fundamentally from the preceding two because both *F* (*q*) and *D*(*q*) are inferred from *nonequilibrium* trajectories. Two cases are considered. In the first (setup 1, blue symbols in Fig. 6b), the analytical form of the free energy is fixed while only the diffusion profile is optimized. This setup represents a situation in which the free energy profile has been independently determined using any of the enhanced sampling techniques to identify the transition state. In the second case (setup 2, red symbols in Figs. 6a and 6b), both *F* (*q*) and *D*(*q*) are optimized simultaneously, corresponding to a situation where only the transition-state location is known in advance, but not the detailed shape of *F* (*q*). In both cases, the true diffusion profile is accurately recovered. The free energy profile is also well reproduced when both *F* (*q*) and *D*(*q*) are optimized. As expected, the second case required substantially more optimization steps, with the precise number depending on the maximum allowed Monte Carlo step size. Overall, this example demonstrates that even a limited set of nonequilibrium trajectories contains sufficient information to reliably extract both *F* (*q*) and *D*(*q*) using our approach.

**Figure 6:**
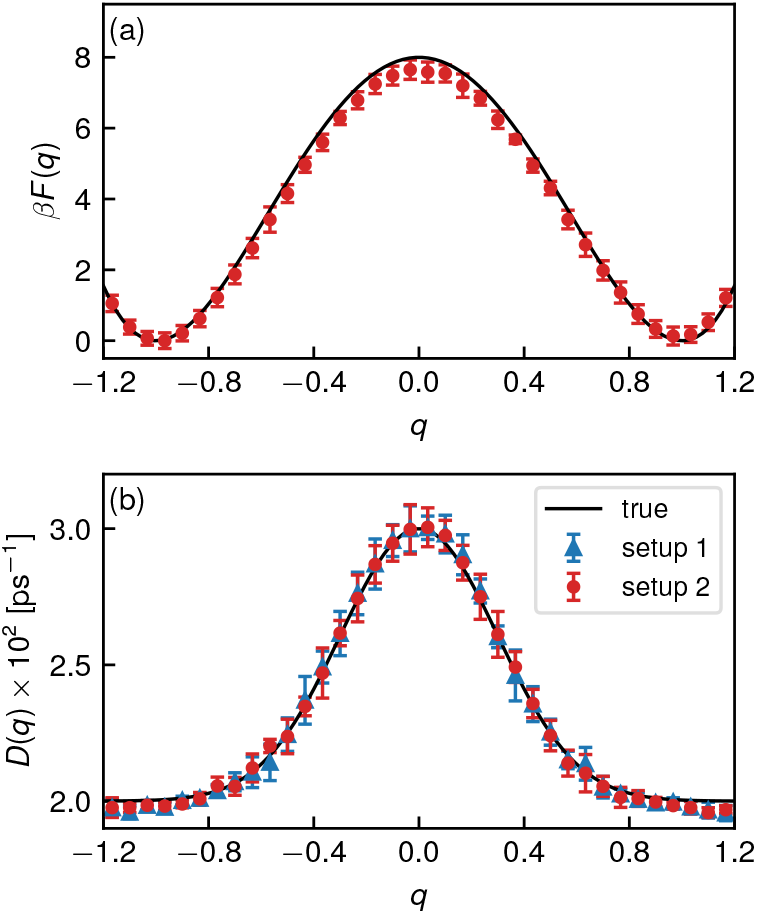
(a) Comparison between the true (black line) and reconstructed free energy profile (red symbols). (b) Comparison between the true diffusion profile (gray line) and the reconstructed diffusion profile obtained by optimizing either only *D*(*q*) (red symbols) or both *F* (*q*) and *D*(*q*) (blue symbols) simultaneously.

### 4 Conclusions

In summary, we have presented a variational method for efficient parameterization of over-damped Langevin models from projected molecular dynamics trajectories in terms of free-energy and diffusion profiles. When a Markovian time series with high temporal resolution is available, the method is particularly efficient because analytical propagators can be employed. For more coarsely sampled trajectories, we proposed a robust but computationally more demanding alternative. A further advantage of the approach is that evaluation of the loss function is computationally inexpensive.

We also emphasized efficient minimization of the loss function and demonstrated how effective Monte Carlo moves can be generated within our framework. Similar speed-ups could be achieved in likelihood-based methods by retaining successful Monte Carlo proposals to guide future updates, although this would require additional tuning of parameters that control how past proposals are replaced by new ones.

Applying the method on three diverse representative systems encompassing both equilibrium and nonequilibrium conditions demonstrate its robustness, efficiency and accuracy. Our aim for future work is to extend the present framework to multidimensional collective-variable spaces.

## Acknowledgements

The authors acknowledge the financial support under Grant Nos. P1-0010 and J1-50033 from the Slovenian Research and Innovation Agency. The use of ChatGPT-5 and Gemini 2.5 Flash for assistance in improving the clarity and readability of the manuscript text is also acknowledged.

## Author Declarations

### Author Contributions

**Anže Hubman**: Conceptualization (equal), Formal analysis (lead); Methodology (equal); Validation (lead); Visualization (lead); Writing – original draft (lead). **Franci Merzel**: Conceptualization (lead); Formal analysis (equal); Funding acquisition (lead); Methodology (equal); Supervision (lead); Validation (equal); Writing – original draft (supporting); Writing – review and editing (equal).

## Data Availability Statement

Data available on request from the authors.

